# Prefrontal control of hippocampal map reorganization during sleep

**DOI:** 10.64898/2026.09.15.751737

**Authors:** Ruoyu Huang, Masahiro Nakano, Claudia Clopath, Jan Born, Thorsten Bartsch, Martin Both, Eduardo Blanco-Hernández, Andrea Burgalossi

## Abstract

During wakefulness, exposure to novel experiences induces hippocampal remapping, in which place cell ensembles reorganize to form distinct representations of new information. Classical models of systems consolidation posit that subsequent sleep stabilizes these wake-formed representations and supports their gradual transfer to medial prefrontal cortex (mPFC) networks primarily via bottom-up hippocampal reactivations. Using miniscope Ca^2+^ imaging of CA1 place cells, we monitored hippocampal activity as mice explored a familiar environment followed by a similar novel one. As expected, place cells remapped in the novel context. However, a single episode of post-encoding sleep selectively reversed remapping dynamics by promoting the reinstatement of familiar place cell representations in the novel environment – an effect that was absent in the wake condition. This neural reinstatement was accompanied by behavioral indicators of familiarity, including reduced exploration and rearing. Chemogenetic inactivation of the mPFC during post-encoding sleep abolished this generalization effect, and instead preserved the novel hippocampal map. Thus, our results extend current models by showing that sleep not only consolidates wake-established hippocampal patterns, but actively shapes hippocampal maps via prefrontal control, thereby supporting memory generalization.

## Introduction

The formation of generalized memories allows organisms to respond adaptively to novel situations by extracting shared regularities across experiences, supporting flexible decision-making and problem-solving^1,2^. Evidence suggests that generalization critically depends upon systems memory consolidation processes, with sleep playing a key role in transforming newly encoded experience into stable and more abstract representations^3–5^. A widely accepted view of systems consolidation posits that during learning, the hippocampus rapidly encodes new information, enabling the acquisition of ongoing experiences while minimizing interference with existing neocortical knowledge^6–8^. During slow-wave sleep, the repeated reactivation of hippocampal place cell ensembles – neurons that fire selectively in specific locations to form cognitive maps^9–12^ – is thought to gradually transfer newly acquired information to cortical networks, including the medial prefrontal cortex (mPFC), thereby promoting the emergence of generalized, schema-like representations^1,2,13,14^. Within this classical framework, hippocampo– mPFC interactions have been largely conceptualized as unidirectional, with hippocampal activity driving cortical reorganization during offline states. While extensive experimental and theoretical work supports this view^3–5,7,10,15^, more recent findings indicate that the mPFC can modulate hippocampal reactivations in a top-down manner during sleep^16^. However, the functional significance of these offline mPFC-to-hippocampal interactions, and whether they actively reshape hippocampal representations formed during learning, has remained unresolved. Hippocampal place cell dynamics have been classically used as a neural readout of spatial generalization and discrimination^17–19^. Depending on the degree of contextual change, place cells may remain stable across contexts, reflecting generalization, or undergo remapping, signaling discrimination between experiences^18,20–22^. Thus, the relative balance between place map stability and remapping provides a quantitative measure of how the hippocampus encodes and differentiates episodic experiences. Importantly, these remapping dynamics (occurring during the wake learning phase) are traditionally attributed to intrinsic hippocampal circuit mechanisms^17^ and have been shown to be stabilized during subsequent sleep^3–5,11^. Whether sleep actively reshapes these hippocampal representations – and thereby contributes to generalization or discrimination – remains an open question.

To address these questions, we combined CA1 place cell recordings with behavioral analysis and chemogenetic manipulation of mPFC inputs. Using a spatial paradigm in which animals explored familiar (F) and overlapping novel (N) environments, we aimed at assessing how the hippocampus resolves conflicts between stored representations and new congruent inputs, as well as the influence of sleep and prefrontal activity on this process.

## Results

### Post-encoding sleep promotes the reinstatement of familiar representations in novel contexts

To monitor hippocampal dynamics during spatial memory processing, we recorded CA1 place cells in freely moving mice using in vivo Ca^2+^ imaging (**Fig. 1a-c** and **Extended Data Fig. 1**) as they ran on U-shaped linear tracks (**Fig. 1a** and **Extended Data Fig. 2**). Prior recordings, animals were habituated to a familiar environment (F) over four consecutive days (**Fig. 1a**; see Methods). During encoding, animals explored the familiar environment (10 min) followed by a novel one (N, 10 min), which differed only in local visuo-tactile cues, while distal contextual cues remained the same (see Methods and **Extended Data Fig. 2**). After a 2-hour retention interval of sleep or wakefulness (**Fig. 1a** and **Extended Data Fig. 3a-c**), animals were re-exposed to the same novel (N’) and familiar (F’) environments (10 min each) in a paired within-subject design. Representative place cell data from one animal are shown in **Fig. 1d**. As expected, N elicited remapping in CA1 place cells (**Fig. 1d**). To quantify changes in spatial representations across environments, we computed standard measures of place map similarity: cell-by-cell rate map correlations, and population vector (PV) correlations (see Methods). Strikingly, post-encoding sleep increased similarity between N’ and F representations, an effect absent in the wake condition (F–N’ cell-by-cell correlations in sleep: 0.45±0.37; in wake: 0.1±0.36; Kruskal-Wallis-test: *p*=5.22e-12, post-hoc Dunn’s test: *p*=1.13e-06. F–N’ PV correlation in sleep: 0.4±0.2; in wake: 0.1±0.11, Kruskal-Wallis-test: *p*=4.22e-16, post-hoc Dunn’s test: *p*=2.21e-06; **Fig. 1e-h**). Thus, this analysis points to the reinstatement of the familiar map in the novel context after sleep, indicative of sleep-dependent generalization.

**Fig. 1:**
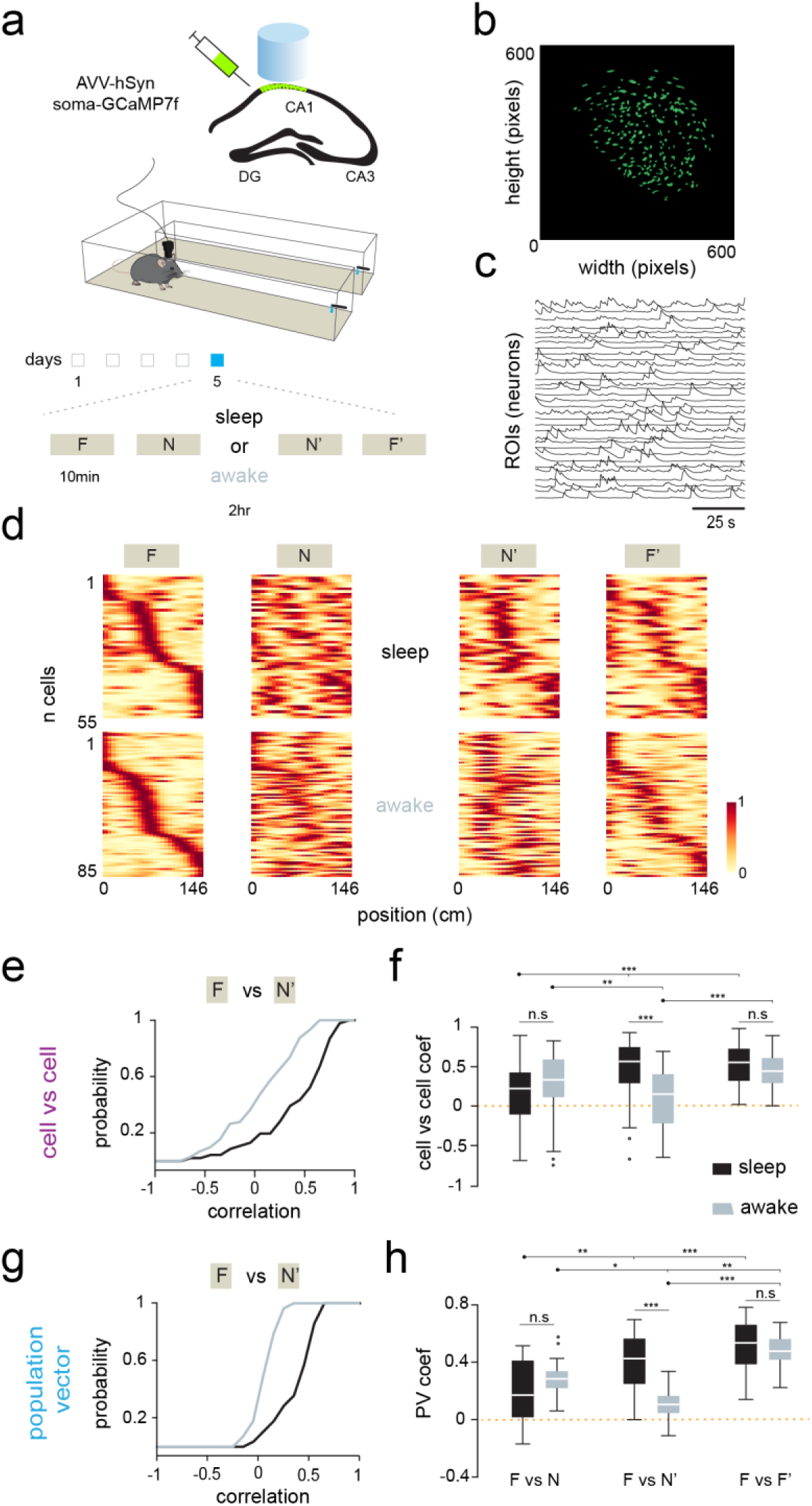
Monitoring CA1 place cell representations with miniscope Ca^2+^ imaging across sleep and wake. (**a**) Schematic of the experimental design. **Top**: Viral injection (green) into dorsal CA1 and GRIN lens implantation for in vivo Ca^2+^ imaging. **Bottom**: U-shaped linear track and behavioral paradigm. Mice were sequentially exposed to a familiar (F) and a novel (N) environment; following a 2-hour retention period of either sleep or wakefulness, they were re-exposed to the same contexts (N’ and F’). (**b**) Representative Ca^2+^ imaging field of view showing neuronal footprints in a representative session (after preprocessing, see Methods and **Extended Data Fig. 1**). (**c**) Example Ca^2+^ traces (quantified by Minian after motion correction and ROI assignment) of randomly selected neurons from (b). (**d**) Linearized place cell rate maps from a representative animal, showing cells co-recorded across all four environments (top: sleep: n_cell_=55; Bottom: wake n_cell_=85). Place maps are aligned to the familiar environment (F). Color bar indicates normalized event rate. (**e**) Cumulative distribution of F-N cell-by-cell correlations from the data shown in (d) in sleep (black) and wake (gray) conditions. (**f**) Boxplot of cell-by-cell correlations across all conditions (data as in (d)) in sleep (black, n_cell_=55) and wake (gray, n_cell_=85) conditions (Kruskal-Wallis test, *p*=5.22e-12; Post-hoc Dunn’s test with Holm-Sidak correction, F-N’ sleep to wake: *p*=1.13e-06). (**g**) Same as in (e) but for population vector (PV) correlations. (**h**) Same as (f) but for PV correlations (black: sleep, gray: wake; n_PV_=28, Kruskal-Wallis test, *p*=4.22e-16; Post-hoc Dunn’s test with Holm-Sidak correction, F-N’ sleep to wake: *p*=2.21e-06).

Altogether, we recorded 8 paired sleep-wake trials from 5 mice (mean sleep durations, 48.99±15.28 min in sleep; 0 min in wake; **Fig. 2** and **Extended Data Fig. 3a-c**). To assess generalization, we measured the similarity between novel post-retention maps (N’) and the initial familiar environment (F) across animals. In line with the example in **Fig. 1d-h**, pooled results (**Fig. 2a-c**) also showed that N’ representations were significantly more similar to F in the sleep than in the wake condition (F–N’ cell-by-cell correlations in sleep: 0.32±0.11, in wake 0.1±0.07. Paired samples t-test: *p*=0.001; F–N’ PV correlations in sleep: 0.31±0.11, in wake: 0.11±0.08. Paired samples t-test: *p*=0.003, n_trials_=8; N_mice_=5; **Fig. 2a,b**; a similar result was observed when including in the analysis all recorded neurons, independent of place-cell classification: F–N’ correlation sleep vs wake, p<0.004; not shown). Sleep did not affect familiar map stability (F–F’ cell-by-cell correlations in sleep: 0.43±0.06, in wake 0.37±0.08. Paired samples t-test: *p*=0.104; F–F’ PV correlations in sleep: 0.42±0.07. In wake 0.37±0.12. Paired samples t-test: *p*=0.14; n_trials_=8; N_mice_=5; **Fig. 2b**). Likewise, mean activity rates and the proportion of active neurons were comparable across sleep and wake conditions (**Extended Data Fig. 4a**). However, following sleep, the proportion of place cells and their within-environment stability in N’ increased toward familiar-like levels (**Extended Data Fig. 4b,c)**, paralleling the reinstatement of familiar representations observed in the correlation analyses in **Fig. 2b**. Notably, although based on a limited number of observations, bootstrap analysis suggested a positive association between sleep duration and F–N’ similarity (cell-by-cell: R^2^=0.59; *p*=0.03; PV: R^2^=0.58; *p*=0.03; n_trials_=8, N_mice_=5; **Extended Data Fig. 3d,e**), consistent with a potential role of sleep in promoting the reinstatement of familiar representations in the novel context. Additionally, a second sleep trial confirmed that F−N’ correlations remained higher after both sleep trials compared to wake (Kruskal-Wallis test: p=0.017, N_mice_=4, n_trials_=4; Bootstrapping of mean differences, 10^4^ repetitions, *p*=0.0004, one-tailed 95% confidence interval; see **Fig. 2c** and **Extended Data Fig. 3g**) indicating the effect was independent of the sleep-wakefulness testing order.

**Fig. 2:**
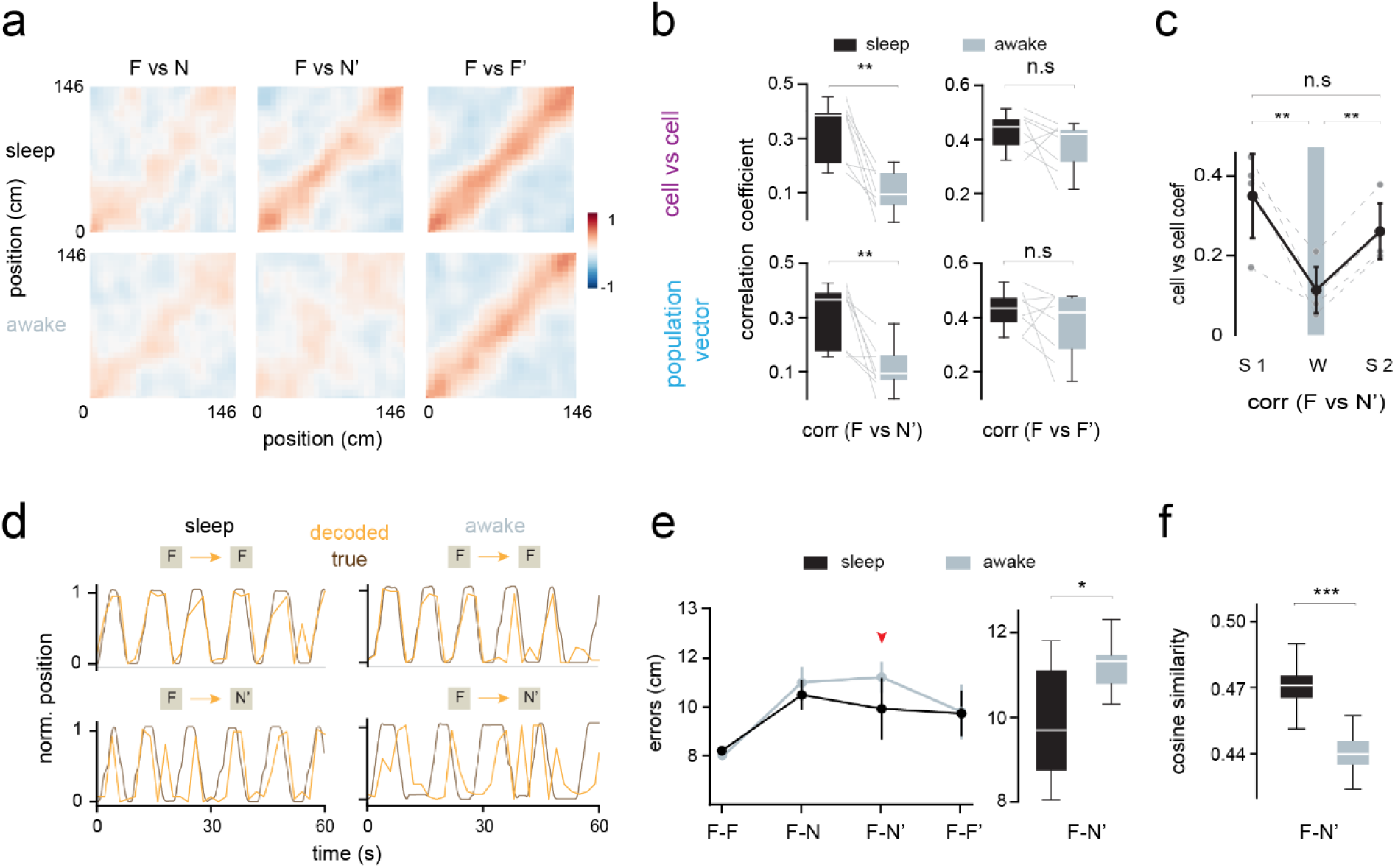
Sleep increases similarity of novel context CA1 place representation to familiar context representation. (**a**) Average spatial correlation matrices between F and subsequent context exposures (N, N’, and F’) in the sleep and wake conditions (n_trials_=8 paired sleep-wake trials; N_mice_=5). Color bar indicates Spearman correlation coefficients. A stronger diagonal band between F-N’ in the sleep condition indicates higher map similarity. (**b**) Boxplots showing pairwise map correlation between F and N’ (left column) and between F and F’ (right column) under sleep (black) and wake (gray) conditions (n_trials_ as in (a)). Both cell-by-cell correlations (top, left) and PV correlations (bottom, left) showed significantly higher F-N’ correlations after sleep, indicating generalization (cell-by-cell F-N’ correlations: *p*=0.001, Paired sample t-test; n_trials_=8, N_mice_=5; PV F–N’ correlations: *p*=0.003, n_trials_=8, N_mice_=5), whereas F-F’ correlations (right) were unaffected (cell-by-cell F–F’ correlations: *p*=0.104, Paired sample t-test; n_trials_=8, N_mice_=5; PV F–F’ correlations: *p*=0.14, n_trials_=8, N_mice_=5). (**c**) Cell-by-cell place map correlation between F and N’ in consecutive sleep-wake-sleep conditions. Generalization was consistently higher in both sleep trials (S1, S2) than wake (W) (Bootstrapping of mean differences, *p*=0.0004, n_trials_=4, N_mice_=4; see Method and **Extended Data Fig. 3g**), indicating the sleep-wake order had no impact on the effect. Error bar indicating mean ± SD. (**d**) Representative traces of predicted position using a maximum likelihood decoder (see Methods). Decoded positions (orange) tracked the true trajectory (brown) less accurately in N’ following wakefulness, whereas decoding accuracy was enhanced after sleep. (**e**) **Left**: Mean absolute decoding error across environments under sleep (black) and awake (gray) conditions. Error bar indicating mean ± SD. **Right**: Boxplot comparison of errors in N’ (arrowhead). Errors in N’ after wakefulness were significantly higher than after sleep (Mann-Whitney U test, *p*=0.018, n_sleep trials_=9, n_wake trials_=8, N_mice_=5). (**f**) Boxplot of unbiased nearest-neighbor population vector similarity between F and N’ based on Ca^2+^ activity from all cross-registered neurons (see Methods). Cosine similarity of Ca^2+^ population vectors was significantly higher after sleep than wake, indicating enhanced reinstatement of familiar population activity patterns during sleep (Mann-Whitney U test, *p*=5.07e-34, n_resample_=100, n_trials_=8, N_mice_=5).

To corroborate these findings, we performed a population decoding analysis to predict the animal’s position from place-cell activity^23,24^. If sleep promotes the reinstatement of the familiar representation in N’, a decoder trained on familiar maps (F) should better predict position in the novel environment (N’) after sleep than wake. Consistent with this, decoding accuracy in N’ was significantly higher following sleep (decoding error in sleep: 9.93±1.23 cm, n_trials_=9; in wake: 11.23±0.64 cm, n_trials_=8; Mann-Whitney U rank test: *p*=0.018; N_mice_=5; **Fig. 2d-e, Extended Data Fig. 3f**). To further assess reinstatement of familiar population activity patterns in N’, we performed a nearest-neighbor analysis of Ca^2+^ population vectors using all cross-registered cells, independent of place-cell classification, spatial location, or temporal order (see Methods). Consistent with previous analyses, F–N’ similarity was higher after sleep than wake (cosine similarity: sleep, 0.47±0.01; wake, 0.44±0.01; Mann–Whitney U test, *p*=5.07e-34; **Fig. 2f**), indicating reinstatement of familiar activity patterns.

Altogether, these data indicate that post-encoding sleep selectively promotes the reinstatement of familiar hippocampal representations in a novel context, indicative of a sleep-dependent shift toward generalization.

### Sleep-dependent neural generalization is reflected in exploratory behavior

To assess generalization behaviorally, we quantified markers of novelty and familiarity, including locomotion speed, stopping events, and rearing^25,26^. As expected, we found that exposure to the novel environment (N) decreased locomotion speed (**Fig. 3a**) and increased stopping frequency (**Fig. 3b**). However, upon re-exposure to the novel environment (N’), sleep and wake conditions diverged: post-sleep animals exhibited higher locomotion speed (normalized speed in sleep: 2.49±0.51; in wake: 1.72±0.66; Wilcoxon signed-rank test: *p*=0.005, n_trials_=12, N_mice_=7; **Fig. 3a**), fewer stopping events (stopping events in sleep: 18.5±11.38; in wake: 37.42±22.04; Wilcoxon signed-rank test: *p*=0.007, n_trials_=12, N_mice_=7; **Fig. 3b**), and reduced rearing (rearing events in sleep: 23.25±10.33; in wake: 44.75±25.08; Wilcoxon signed-rank test: *p*=0.01, n_trials_=12, N_mice_=7; **Fig. 3c**), resembling behavior in F. Importantly, the correlation between rearing and decoding error between sleep and wake conditions was significantly different (Regression coefficient permutation test, 10^4^ repetitions, ΔR^2^: *p*=0.018; Δslope: *p*=0.012, one-tailed 95% confidence interval; n_trials_=8, N_mice_=5; **Fig. 3c, Extended Data Fig. 3h**) with the correlation in sleep condition showing a positive trend (R^2^=0.15, *p*=0.34), suggesting a sleep-dependent link between hippocampal representations and behavior.

**Fig. 3:**
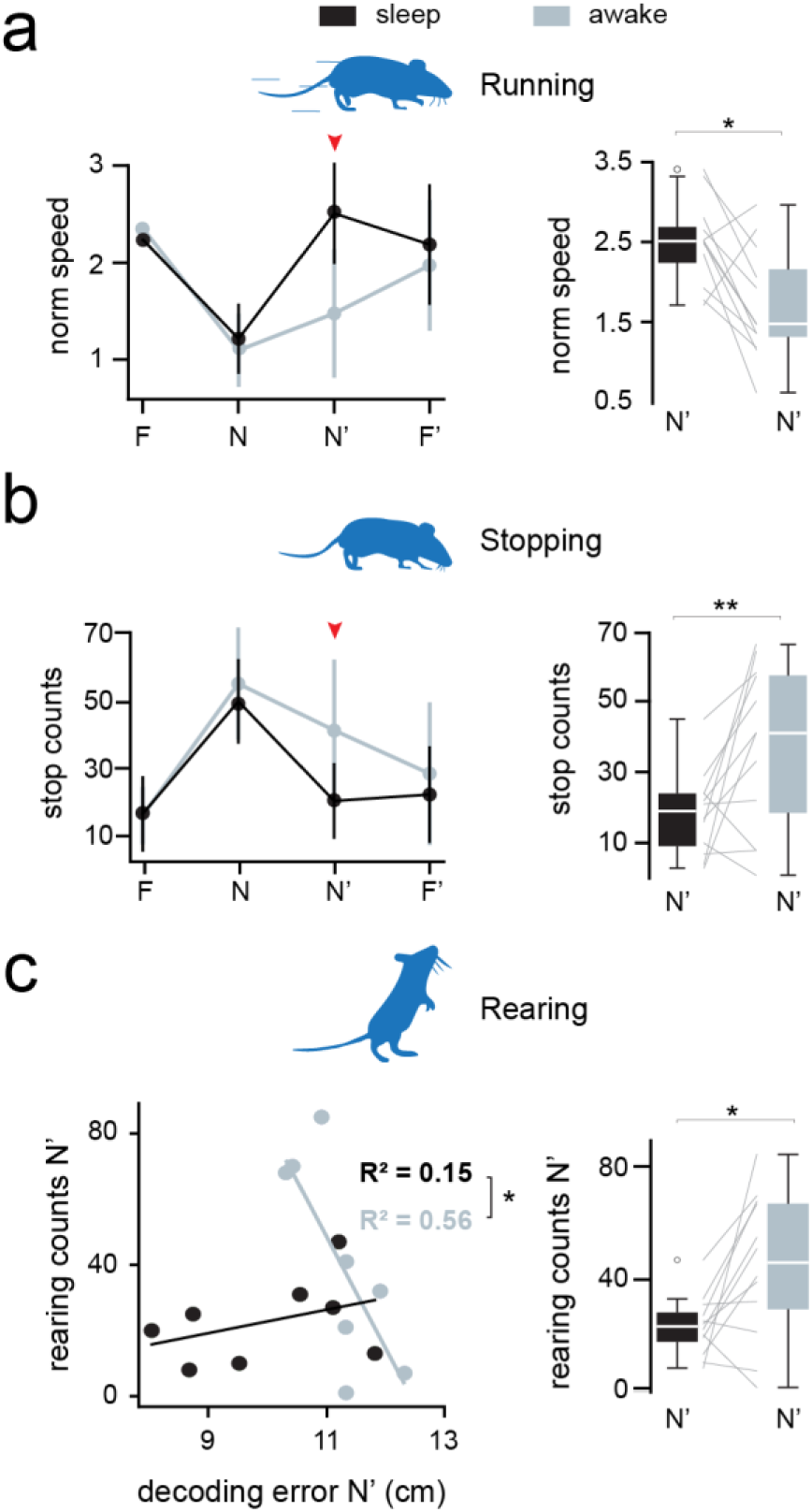
Sleep-mediated reinstatement of familiar place representations is accompanied by behavioral indicators of familiarity. (**a**) **Left**: Normalized locomotion speed across environments for the sleep (black) and wake (gray) conditions (n_trials_=12, N_mice_=7). Speed was calculated in the linear arms, outside of reward locations (see Methods). Error bar indicating mean ± SD. **Right**: Boxplot comparison of velocities in N’ (arrowhead). Speed was significantly lower in N’ after wake than after sleep (Wilcoxon signed-rank test, *p*=0.005, n_trials_=12, N_mice_=7). (**b**) **Left**: Stopping events across environments under sleep (black) and wake (gray) conditions (n_trials_=12, N_mice_=7). Stops were defined as phase with locomotion < 1 cm/s outside reward zones (see Methods). Error bar indicating mean ± SD. **Right**: Boxplot comparison of stopping events in N’ (arrowhead). Number of stopping events was significantly higher in N’ after wake than after sleep (Wilcoxon signed-rank test, *p*=0.007, n_trials_=12, N_mice_=7). (**c**) **Left**: Spontaneous rearing events in N’ versus decoding error for sleep (black) and wake (gray) conditions (n_trials_=8, N_mice_=5). The correlation between rearing and decoding error between sleep and wake conditions was significantly different (Regression coefficient permutation test, 10^4^ repetitions, ΔR^2^: *p*=0.018; Δslope: *p*=0.012; see **Extended Data Fig. 3h**). **Right**: Rearing counts was significantly higher in N’ after wake than after sleep (Wilcoxon signed-rank test, *p*=0.01, n_trials_=12, N_mice_=7), indicative of behavioral generalization and consistent with data in (a) and (b).

### Prefrontal cortex activity is required for sleep-dependent hippocampal generalization

Repeated experiences across similar environments have been shown to support the formation of generalized “schema-like” representations in the mPFC^27,28^. In our task, repeated pre-exposure to the F environment likely allowed the mPFC to form a generalized representation of the task structure. Hence, we hypothesized that sleep might engage mPFC–hippocampal interactions that support the generalization observed in our data (**Fig. 4**). To test this hypothesis, we bilaterally expressed the inhibitory DREADD hM4Di in the mPFC and chemogenetically silenced mPFC neurons by administering deschloroclozapine (DCZ^29^; i.p.) before sleep (as control, vehicle injections were performed in a paired within-subject design; n_trials_=4, N_mice_=4; **Fig. 4a-c**). Both control and vehicle conditions showed comparable sleep during the retention period (sleep duration 63.02±13.63 min in DCZ; 60.76±24.65 min in vehicle, **Extended Data Fig. 3c**). However, only mPFC inactivation reduced the reinstatement of the familiar place-cell representation in N’ (**Fig. 4d**) and increased the stability of the novel representation (**Fig. 4e**), pointing to a shift in hippocampal dynamics from generalization toward discrimination. To quantify this effect, we computed a “Generalization Index” defined as the distance from the identity line relating F–N’ and N–N’ similarities for each data point (**Fig 4f,g**). Indeed, the mPFC inactivation group exhibited a significantly reduced Generalization Index, indicative of a bias toward consolidation of the N’ map relative to both wake and intact-sleep control conditions (Generalization Indices: in DCZ: −0.08±0.05, n_trials_=4; in sleep control: 0.09±0.04, n_trials_=12; in awake: −0.05±0.05, n_trials_=8; Kruskal-Wallis test *p*=0.0002; **Fig. 4g**). Together, these results indicate that mPFC activity during sleep is required to bias hippocampal representations toward previously established familiar maps, revealing a causal top-down mechanism by which cortical activity shapes hippocampal memory reorganization and supports generalization.

**Fig. 4:**
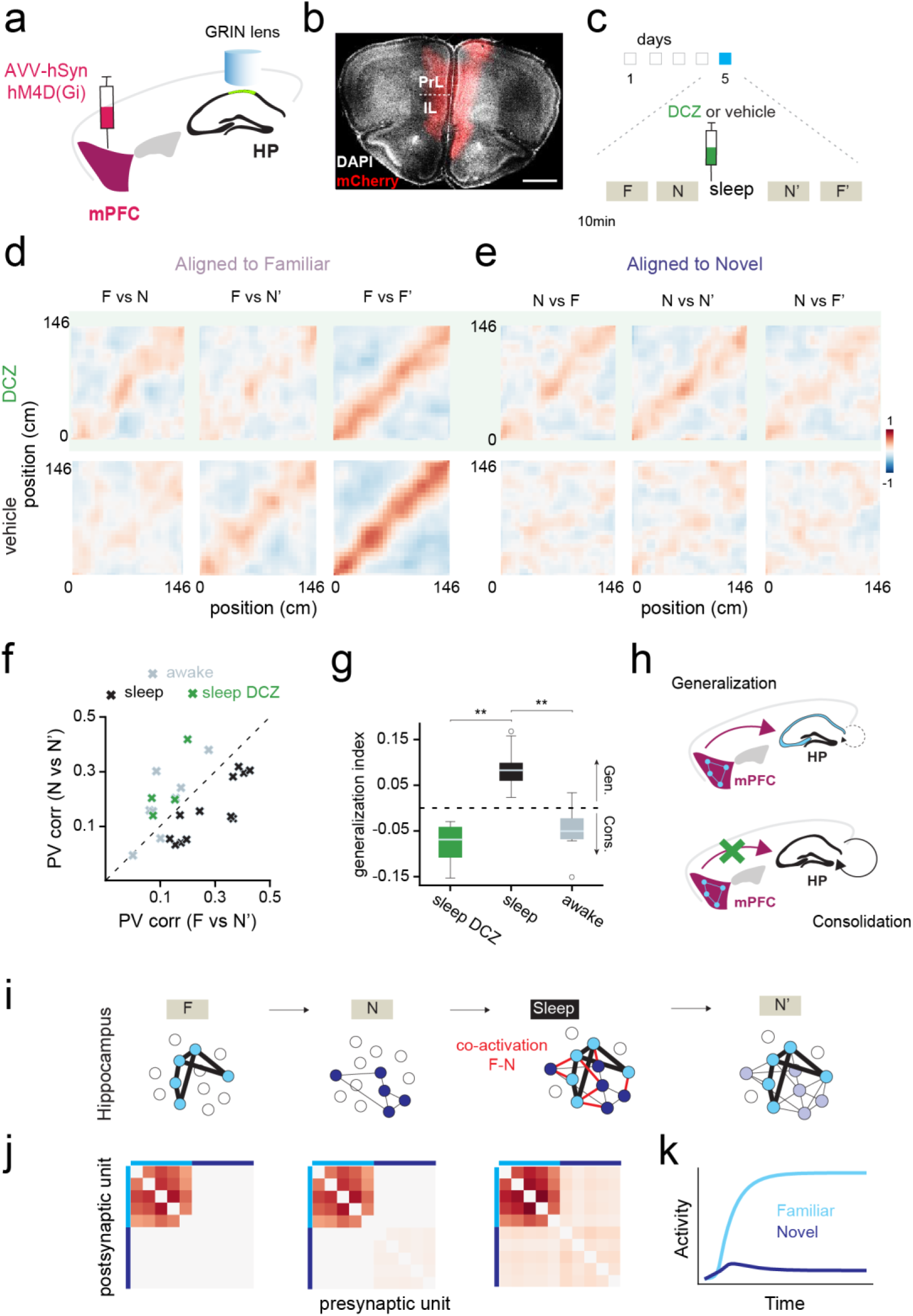
mPFC activity during sleep supports generalization in CA1 representation. (**a**) Schematic of the experimental design for the miniscope Ca^2+^ recording in CA1 and chemogenetic inactivation of the mPFC via bilateral expression of inhibitory DREADD hSyn-hM4D(Gi)-mCherry. (**b**) Representative brain section showing expression of hM4D(Gi)-mCherry in mPFC in both hemispheres. Red: mCherry; Gray: DAPI. Scale bar: 1 mm. PrL: prelimbic cortex, IL: infralimbic cortex. (**c)** Schematic of the behavioral paradigm. Same as in Fig. 1a, with the difference that prior to the retention phase, mice received an intraperitoneal injection of either deschloroclozapine (DCZ, 0.1mg/kg) or vehicle. (**d**) Average spatial correlation matrices between F and subsequent environments (N, N’, and F’) under the DCZ and vehicle conditions (n_trials_=4 paired trials; N_mice_=4). Note the high F–N’ similarity in the control (vehicle) condition, indicative of generalization, and the reduced similarity in the DCZ condition. Color bar indicates Spearman correlation coefficients. (**e**) Same data as in (d), but aligned to the N environment. Note the high N–N’ similarity observed under DCZ, indicating enhanced consolidation of novelty during mPFC inhibition. (**f**) Scatter plot of population vector (PV) correlations corr(F,N’) vs corr(N,N’) showing representation bias across conditions. Each point represents the mean correlation coefficient of one trial (DCZ: n_trials_=4, N_mice_=4; Sleep and vehicle control: n_trials_=12, N_mice_=9; Awake: n_trials_=8, N_mice_=5). The dashed line indicates equal similarity to F and N; points above indicate consolidation toward N, whereas points below indicate generalization toward F. (**g**) Quantification of the data shown in d–f using a Generalization Index, which measures the representational bias in N’ based on the signed distance of each data point in (f) from the identity line (see Methods). Positive values indicate generalization toward F, whereas negative values indicate consolidation toward N. Note that sleep (black) shifted CA1 representations toward generalization, whereas mPFC inhibition (green) shifted them toward consolidation. (Kruskal-Wallis test, *p*=0.002; Post-hoc Dunn’s test with Holm-Sidak correction, *p*_(DCZ vs sleep)_=0.003, *p*_(sleep vs awake)_=0.002, *p*_(DCZ vs awake)_=0.644). (**h**) Schematic summarizing the experimental findings. Top: During sleep, top-down mPFC– hippocampal interactions promote reinstatement of familiar representations in CA1. Bottom: mPFC inactivation during sleep shifts the balance away from generalization, biasing CA1 representations toward consolidation of the novel (N) representation. (**i)** Schematic model of mPFC–hippocampal interactions underlying sleep-dependent generalization. Behavioral exposure establishes strong connectivity within F (light blue) and weak connectivity within N (dark blue) hippocampal representations. During sleep, F–N co-activation enabling subsequent novel inputs to recruit familiar representations (shown by the red lines) and shift network dynamics toward a generalized map. (**j)** Visualization of the weight matrix after experiencing F (left), N (middle), and sleep (right). The color bar labels indicate the F and N neurons. Darker red indicates a stronger connection. (**k)** Average activity of the F and N representations in N’ after sleep.

To gain mechanistic insight, we developed a simple computational model comprising hippocampal populations representing F and N environments, with recurrent connectivity shaped by Hebbian plasticity and stronger in F due to greater experience (**Fig. 4h-k**). Because the nature of top-down mPFC signals during sleep remains unknown, we did not explicitly model prefrontal inputs. Instead, motivated by evidence that mPFC activity coordinates hippocampal reactivations^16,30–32^, we asked whether co-reactivation of F and N representations alone could promote generalization. In the model, co-activation of F and N during the sleep phase strengthened cross-population connections through Hebbian plasticity (**Fig. 4i,j**). Following this sleep-dependent coupling, sensory input in N’ initially activated the N population, which in turn recruited the F population through the newly formed connections. Because F exhibited stronger recurrent connectivity and competed under global inhibitory normalization, network activity was biased toward the F representation, rendering the N’ response more similar to F (**Fig. 4k**). Thus, with minimal assumptions, the model shows that sleep-dependent co-reactivation of F and N representations can be sufficient to bias hippocampal consolidation toward generalization.

## Discussion

Our findings reveal a previously unrecognized role of sleep in actively reshaping hippocampal representations through top-down prefrontal input, providing a mechanistic link between cortical control and hippocampal memory generalization. Classical models of systems consolidation have largely emphasized hippocampal-prefrontal interactions, during which hippocampal reactivations stabilize wake-formed representations and gradually integrate them into cortical networks^33,34^. Within this bottom-up framework, hippocampal maps formed during wake are assumed to be consolidated with minimal modification. Our data extend this view by showing that post-encoding sleep can selectively reverse wake-induced hippocampal remapping, and actively bias hippocampal representations toward generalization.

Consistent with prior work^17,27,35–40^, we found that altering local sensory cues was sufficient to induce robust reorganization of CA1 place fields (**Fig. 1**, **Fig. 2 and Extended Data Fig. 4**). Such remapping is typically attributed to changes in sensory inputs^17,41–43^ and is largely thought to arise from intrinsic hippocampal circuit mechanisms. Here, we show that a single episode of post-encoding sleep can counteract encoding-related remapping and bias hippocampal representations toward a pre-existing map. Specifically, sleep promoted the reinstatement of familiar place cell ensembles in the novel context (N’), increasing their similarity to the original (F) representation. Remarkably, chemogenetic inactivation of the mPFC during post-encoding sleep abolished the reinstatement of familiar representations. Thus, these findings indicate that hippocampal remapping dynamics are more flexible and sleep-sensitive than previously recognized, and are not exclusively determined by intrinsic hippocampal circuitry. Rather, remapping established during wake can be reorganized during sleep under the influence of mPFC input, highlighting a prefrontal contribution to hippocampal remapping dynamics and to the emergence of generalized spatial representations.

Notably, while the novel hippocampal map (N) was not consolidated under normal sleep or wake conditions (**Fig. 1-3**), mPFC inactivation during sleep allowed the novel representation to stabilize (**Fig. 4**). This suggests that, in the absence of prefrontal input, hippocampal maps follow a default consolidation process consistent with classical complementary learning systems model^15,44^. However, when prior congruent representations are present in prefrontal circuits, mPFC activity during sleep can promote the integration of novel hippocampal representations into existing schemas. Together, these findings point to a competitive prefrontal–hippocampal interaction during sleep that regulates whether novel representations are stabilized independently or assimilated into generalized representations^45^. Anatomically, the mPFC and hippocampus are connected indirectly via the nucleus reuniens and perirhinal/entorhinal pathways^7,46–48^, while functional studies have provided compelling evidence for their bi-directional interaction during sleep^16,49,50^ involving both excitatory and inhibitory influences^48,51–53^. Because the nature of the instructive signal provided by the mPFC during sleep remains unknown, we did not explicitly model prefrontal input. Instead, we focused on biologically plausible intra-hippocampal mechanisms through which coordinated reactivation of existing and novel representations could support generalization. Motivated by evidence that prefrontal activity coordinates hippocampal reactivation during sleep^16,30–32,50^, we implemented a minimal mechanism in which transient co-activation of F and N representations promotes Hebbian strengthening of cross-population connections. Such hippocampal linking mechanism could account for the observed reinstatement of the familiar representation in N’ (**Fig. 1,2**) and yields a directly testable prediction for future work.

Extensive work in cognitive neuroscience has established that prefrontal–hippocampal interactions are central to episodic memory retrieval, particularly in models of schema-guided recall and reconstructive memory, where prior knowledge systematically shapes the reconstruction of past experiences^1–7^. More recently, evidence from both humans and rodents has extended this framework to encoding, suggesting that prefrontal schemas can influence the assimilation of new episodic information^14,54–56^. However, whether such top-down interactions also operate during offline states, and specifically during sleep, has not been established. Here we show that even a brief post-encoding sleep interval (ca. two hours) is sufficient to reshape hippocampal representations and behavior in a manner consistent with “schema-driven” generalization. In this window, hippocampal activity is biased toward the reinstatement of pre-existing representations, resulting in the transformation of novel inputs into more generalized codes, accompanied by corresponding changes in behavior. These findings provide evidence for a previously uncharacterized sleep-dependent mechanism through which top-down mPFC– hippocampal interactions can actively reorganize memory representations to support generalization.

Central to the observed generalization in our data was the high degree of congruency between the F and N environments. Although sufficiently distinct to elicit remapping (consistent with previous work^37,39,40,42,43^), they shared key structural features, including distal cues, arena geometry, and behavioral contingencies. In addition, novelty in our paradigm was of limited behavioral salience, which likely further favored generalization over discrimination. This neural effect was paralleled at the behavioral level, where animals exhibited indices of familiarity in N’, including higher running speed and reduced rearing (**Fig. 3**), resulting in more efficient water collection, consistent with reduced impact of novelty-related exploration. We propose that under these conditions, top-down mPFC influences during sleep bias hippocampal processing away from encoding of novel details and toward reuse of existing representations, thereby supporting an adaptive role of sleep for optimizing behavior in novel (but similar) contexts.

In summary, our findings extend classical systems consolidation models by suggesting that sleep regulates the balance between generalization and discrimination rather than merely stabilizing hippocampal representations. This balance is shaped by top-down mPFC– hippocampal interactions during sleep, which determine whether novel information is integrated into existing schemas or maintained as distinct representations. External modulation of these processes (e.g., via targeted memory reactivation^57,58^ or neurostimulation^59–61^) could offer a potential avenue to influence maladaptive memory processing in disorders characterized by altered balance between generalization and discrimination, such as Alzheimer’s disease^62^, schizophrenia^63^ and post-traumatic stress disorder^64–67^.

## Acknowledgements

We thank Fabio Monteiro for excellent technical assistance and Arya Hassanali for input on data analysis. We thank Katja Lankisch and Matthias Klumpp (University of Heidelberg) for their support with the implementation of miniscope recordings. We thank Niels Niethard for feedback on earlier versions of the manuscript. We also thank Olga Garaschuk (University of Tübingen) for her advice and support regarding viral constructs and Ca^2+^ imaging. This work was supported by the Eberhard Karls University of Tübingen, and the German Research Foundation (DFG grant BU 3126/3-1, DFG grant BO 3512/2-1),

## Author contributions

A.B., J.B., T.B. and E.B-H. conceived and designed the experiments. A.B. and E.B-H. supervised the experiments. M.B. assisted in setting up of Ca^2+^ miniscope recordings. R.H. established the setup and performed the experiments. R.H. and E.B-H. analyzed the data. M.N. and C.C. implemented the model. R.H., J.B., E.B-H. and A.B. wrote the original manuscript. All authors revised and edited the manuscript.

## Data and code availability

All data needed to evaluate the conclusions of the manuscript are presented in the paper and/or supplementary materials, and will be available from a public repository upon publication.

## Methods

### Experimental subjects

Experimental procedures were performed according to German guidelines on animal welfare under the supervision of the local ethics committees (Regierungspräsidium Tübingen, licenses CIN03/20G, CIN1/23G, CIN03/23G). 11 adult male wild-type C57BL/6J mice (>9 weeks old; Charles River, RRID: IMSR_JAX:000664) were used in the present study.

### Surgery and miniscope Ca^2+^ Imaging

General anesthesia and surgical procedures were performed as previously described ^68–70^. Miniscope implantation and registration followed established protocols in ^71–73^. Specifically, mice were anesthetized via intraperitoneal injection of a fentanyl, midazolam and medetomidine mixture and secured in a stereotaxic frame.

Craniotomies were made at AP −1.85 mm, ML +1.25 mm for targeting the dorsal CA1. AAV1-hSyn-somaGCaMP7f (Viral Vector Facility UZH) was injected at DV −1.1 mm using a Hamilton syringe (8.1 x 10E12 vg/mL, 100 nL/min; 500 nL total) or coated on the surface of the GRIN lens, according to published procedures^74^. After injection, the overlying cortex was aspirated using a 30G blunt needle under constant saline irrigation. A 1 mm GRIN (gradient refractive index) lens (GRINTECH, NEM-100-20-20-520-S-0.5p; Klumpp et al. 2025), was implanted above the CA1 and secured with cyanoacrylate and dental cement. In a subset of experiments, bilateral injections of a AAV9-hSyn1-hM4D(Gi)-mCherry (8.8 x 10E12 vg/mL, 70 nL/min; 300 nL; Viral Vector Facility UZH) were targeted to the mPFC (AP +2.0 mm, ML ±1.0 mm, DV −2.6 mm; i.e. prelimbic and infralimbic cortices) using a 17° angle to avoid damage to the superior sagittal sinus, as previously described^76^. Two to three weeks after the surgery, mice were briefly anesthetized with isoflurane to assess GCaMP expression and attach the miniscope baseplate.

Single-photon Ca^2+^ imaging was performed using a UCLA Miniscope V4.4 (OpenEphys)^77,78^. To motivate running, animal received water rewords (or small food pellets) at each end of the track (see section “Behavioral task design and data collection”). Prior recordings, mice were habituated to handling and to a 3D-printed dummy miniscope. Imaging was conducted in CA1 at 20 Hz during four 10-min exploration. The field of view was 608 × 608 pixels (∼1 mm diameter). A commutator (Open Ephys) was integrated into the setup to prevent cable twisting and to counterbalance the miniscope weight during free movement.

### Histology

After experiments, mice were euthanized with an overdose of pentobarbital and perfused with 0.1 M phosphate-buffered saline (PBS) followed by 4% paraformaldehyde (PFA, Sigma-Aldrich). Brains were post-fixed in PFA and sectioned into 70 µm coronal slices using a vibratome. Sections were stained with DAPI (Sigma Aldrich) and imaged with a Zeiss Axio epifluorescence microscope to verify GCaMP7f expression and GRIN lens placement. All animals included in the study showed robust GCaMP7f expression within the CA1 pyramidal layer, and correct placement of the GRIN lens above the CA1 (see representative data in **Fig. 1b** and **Extended Data Fig. 1**).

### Behavioral Task Design and Data Collection

Experiments took place in a 1 × 1 × 1 m sound-attenuated Faraday cage illuminated with dim visible light (21 lux) and infrared light (840 nm). Animal position was tracked at 30 Hz using a top-mounted CMOS camera (DMK33UX265, The Imaging Source) with an IR filter (FELH0800, Thorlabs). A second CMOS camera (DMK23UP1300, The Imaging Source) recorded sleep behavior at 20 Hz. Calcium imaging and behavioral data were synchronized and acquired with the Syntalos system^75^.

Mice were trained on a water-collection task on a transparent U-shaped linear track (arms 60 cm, curve radius 11.5 cm, corridor width 7 cm, total end-to-end distance ∼156 cm), enclosed by 20 cm walls. Water ports (18G blunt needle-tip) equipped with motors (RP-Q1, Takasago Fluidic Systems) and IR sensors (880 nm, MH Series) were positioned at both ends. When a mouse crossed an IR beam, a droplet of water was delivered at the opposite port, requiring mouse to alternate between ends to collect rewards.

Visual and tactile landmarks were included on the walls and floor to support spatial recognition (see **Extended Data Fig. 2**). Five environments were used, differing in local cues (i.e. textures, materials, and colors). However, aspects such as track geometry, water port positions, roof texture, room distal cues and the experimenter remained constant.

Each mouse was assigned one familiar (F) environment and four distinct novel (N) environments (see **Extended Data Fig. 2** and **Fig. 1a**). Mice were pre-exposed to the familiar environment across four days prior to recoding (4 exposure × 10 minute /day). Mice were allowed to sleep during habituation. On recording days, mice first explored the F and then a N environment. This was followed by a 2-hour retention phase under either sleep or wake conditions, followed by re-exposure to the same N’ and F’ environments (see experimental timeline in **Fig. 1a**). For the mPFC inactivation experiments (**Fig. 4**), animals received an intraperitoneal injection of either deschloroclozapine (DCZ; 0.1 mg/kg, dissolved in 2% DMSO in saline^29^) or vehicle/saline solution (control condition) immediately prior to the sleep period. In the sleep condition, mice were returned to their home cage and slept (**Extended Data Fig. 3a-c**). In the wake condition, mice were placed in the same home cage, and wakefulness was ensured by providing novel nesting material, following the procedures described by e.g. ^79^ with gentle handling, if required. During both conditions, the mouse behavior was monitored with overhead camera recording under 44 lux illumination.

In this study, an environment refers to one of the four consecutive 10 min recordings (F, N, N’, or F’), whereas a trial denotes the complete F-N-sleep/wake-N’-F’ experimental sequence. The dataset of neural recordings in **Fig. 2** consisted of 8 paired sleep-wake trials and one unpaired sleep trial from 5 mice (n_trials_=8, N_mice_=5; **Fig. 2**); in **Fig. 4**, it consisted of 4 paired DCZ-vehicle trials from 4 mice (n_trials_=8, N_mice_=4 out of 5 mice; one mouse was excluded from the analysis due to suboptimal viral expression; **Fig. 4**). The additional behavior-only dataset **(Fig. 3**) was performed in two wildtype animals without neuronal recordings and contributed four paired sleep-wake trials (total n_trials_=4, N_mice_=2; **Fig. 3**). For the dataset in **Fig. 2**, each mouse underwent both sleep and wake conditions in a paired within-subject design; three of the five animals contributed two paired sleep-wake datasets each, while one animal was tested twice under sleep and only once under wakefulness, and one animal contributed only a single paired sleep-wake dataset. The corresponding unpaired sleep recording was only included in control analyses for (i) estimating effect sizes in the sleep-wake-sleep (S1–W–S2) condition order (**Fig. 2f**) and (ii) comparing the decoder error in N’ between sleep and wake (**Fig. 2e**). For all other analyses, only paired data were included. For the dataset in **Fig. 4**, each mouse underwent both DCZ and vehicle conditions in a paired within-subject design; each mouse contributed one paired DCZ-vehicle dataset each (n_trials_=8, N_mice_=4). The same conclusions were supported by per-mouse analyses, excluding the possibility of a mouse-specific bias (**Fig. 2**: cell-by-cell F– N’ correlations: *p*=0.002, PV F–N’ correlations: *p*=0.002; **Fig. 4**: Kruskal-Wallis test, *p*=0.002; Post-hoc Dunn’s test with Holm-Sidak correction, *p*_(DCZ vs sleep)_=0.006, *p*_(sleep vs awake)_=0.016, *p*_(DCZ vs awake)_=0.577; not shown)

### Analysis of behavioral data

The animal position during the sampling of the familiar and novel environments was tracked using a custom OpenCV-based Python script. Position data were downsampled to 20 Hz and smoothed using an 800 ms sliding window. The U-shaped track was linearized to a one-dimensional representation.

During exploration, locomotion velocity was computed in the straight segments of the U-shaped arenas as follows: the linearized track was first divided into 30 spatial bins (5.2 cm/bin), and the bins within the curvature and reward zones were excluded. Velocity was averaged within each bin and normalized to the mean velocity from the familiar (F) environment of the same recording day. Stopping behavior was defined as episodes in which animal absolute locomotion velocity fell below 1 cm/s (reward zones excluded). Each stop was quantified as periods when the animal’s velocity dropped below 1 cm/s until it rose above 1 cm/s again. Stop episodes from post-encoding novel environments (N’) were then extracted, and frames containing rearing behavior were manually annotated (**Fig. 3c**).

Sleep episodes during the post-encoding retention phase were quantified from videography (**Extended Data Fig. 3a-c**) following previously published procedures ^80,81^. Motion energy was extracted from a ROI covering the animal cage and analyzed in Python using Facemap (https://github.com/MouseLand/facemap v1.0.6) ^82,83^. Motion traces were smoothed with a 5-second-long median filter, and immobility was defined using a threshold of ∼0.1±0.05 standard deviation of the smoothed motion trace. Classification was verified by manual inspection of body posture in the raw video and the threshold was adjusted within the mentioned range. Episodes of immobility lacking the characteristic sleep posture or shorter than 60 s were excluded. The total sleep duration was computed as the sum of all visually-validated sleep epochs within the retention phase.

### Analysis of Ca^2+^ Activity Data

Ca^2+^ imaging data were preprocessed using Minian-based pipelines^84^, including denoising, motion correction and CNMF-E-based signal extraction^85,86^. Cross-session registration was performed based on centroid distance of cell footprints, and only cells detected in all environments of a recording day were retained.

A custom Python script detected Ca^2+^ events from processed traces. Ca^2+^ traces were aligned with behavioral data, downsampled to 20 Hz, and baseline-corrected using a 20 s median filter. A 2 Hz low-pass filter was then applied. Event candidates were identified as peaks exceeding 2 median absolute deviations (MAD)^87^. To account for somaGCaMP7f indicator decay kinetics^88^, event candidates within 1 second of each other were further filtered: only peaks 4 MAD above the prior and 2 MAD above the following were retained; otherwise, only the first peak was counted.

### Place cell analysis

The linearized tack was divided into 30 bins (5.2 cm/bin). Since CA1 place cells are directional in linear tracks^71^, Ca^2+^ events were separated by animal’s running direction and lap directions were analyzed separately as in^71^. Immobility phases (locomotion speed < 4 cm/s) and the reward zones (first and last bins) were excluded. Spatial rate map per direction were computed as event count per bin divided by occupancy, with occupancy less than 200 ms set to zero. Maps were smoothed with a Gaussian kernel (*σ*=5.2 cm, unless otherwise indicated) and normalized to the maximum value. Neurons at the edge of the GRIN lens or those with few than 5 events per lap direction were excluded from the analysis.

Place fields were defined via a shuffled peak-rate method^89^. For each cell, 500 random-shuffled traces (time-shifted >5 second) were generated. Cells in which the original peak rate exceeded the 95th percentile of shuffled peak rates were classified as place cells.

To assess activity pattern of the same cell across environments, place cells identified in F environment was aligned with their registered matches in the comparison environment. The Spearman correction of each corresponded cell-pair was computed. Then, to assess the similarity of spatial representations at the population level, we computed spatial rate map population vector (PV) correlations across environments, as previously described^24^. Specifically, at each spatial bin, we constructed a PV consisting of the firing rates of all aligned cells. Spearman correlations were computed between PVs at corresponding bins across environments, yielding a correlation profile for each position of the track. For spatial correlation analysis (as shown in **Fig. 1d-h**), only place cells that were consistently identified across all four environments and exhibited F–F’ correlations > 0 were included, while cells that could not be matched across all four environments were excluded.

To quantify whether CA1 spatial representations in the post-retention re-exposure (N’) were more similar to the F or N environment, we computed a Generalization Index defined as:

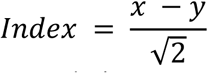

where *x* and *y* represent the rate-map population vector correlations between F and N’, and between N and N’, respectively. The Generalization Index corresponds to the signed perpendicular distance from the identity line (*y* = *x*) in the correlation space (range: -√2 to √2), with 0 indicating no bias, positive values indicating generalization toward F, and negative values indicating consolidation toward N (**Fig. 4g**).

### Maximal Likelihood Decoder

To decode the animal’s position from Ca^2+^ activity data, a Maximum Likelihood Estimation (MLE) decoder^23,24,90^ was trained on the first 75% of Ca^2+^ data in the F (only place cells were included) and tested it on the remaining 25%, as well as on data from other environments. Only place cells that were consistently identified across all four environments and exhibited F–F’ correlations > 0 were included, while cells that could not be matched across all four environments were excluded.

Ca^2+^ events were assumed to follow a Poisson process and to be independent across neurons^90^. Event traces were binned into 2 seconds windows to ensure sufficient temporal resolution for the decoding (**Extended Data Fig. 3f**). For neuron *i* at position *x*, the probability of observing Ca^2+^ event rate *r_i_* can be given by the Poisson probability mass function (PMF)^23,90^, where *λ_i,x_* is the expected non-smoothed event rate at *x* from the training map:

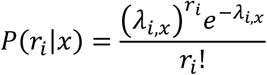

Thus, the probability of observing the population activity *r_t_* of *N* neurons in current time window *t* given that the animal was at position *x* can be computed using a Poisson likelihood model:

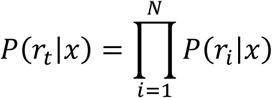

In practice, we computed the log-likelihood to avoid numerical underflow and added a small number to avoid log 0:

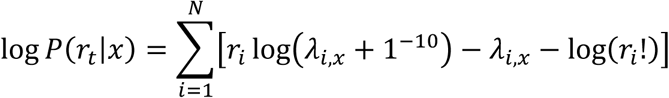

The position *x* that maximized the likelihood was taken as the decoder predicted position for the time window *t*:

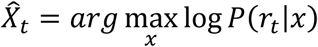

Decoder performance was assessed using mean absolute error (MAE) between decoder predicted position *X̂_t_* and actual positions *X_t_* over *T* time windows in the last step:

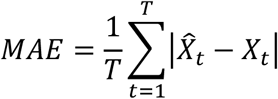

### Nearest-neighbor similarity of population Ca^2+^ activity patterns

To quantify the reinstatement of population activity patterns independent of spatial location and temporal order, we performed a nearest-neighbor matching analysis on population Ca^2+^ traces (**Fig. 2f**). Cross-registered neurons from all animals were pooled, and the number of neurons was equalized to the median by randomly duplicating or removing neurons. Ca^2+^ events occurring during runs outside the reward zones were identified, and the amplitudes of the Ca^2+^ event were convolved with an asymmetric Gaussian kernel (*σ*_rise_=100 ms, *σ*_decay_=750 ms)^40^. The resulting traces were averaged within consecutive 200 ms time bins. For each iteration, 30% of the neurons were randomly sampled without replacement while preserving neuron identity across environments. Each time bin was represented by a population vector of Ca^2+^ activity (Ca^2+^-PV). For every Ca^2+^-PV in F, cosine similarity was computed with all Ca^2+^-PV in N’, and the maximum cosine similarity (nearest-neighbor similarity) was retained. The mean nearest-neighbor similarity across all Ca^2+^-PV in F was used as the similarity score. This procedure was repeated 100 times with independent neuronal subsamples.

### Hippocampal network model

We constructed a simple rate based recurrent network model consisting of 10 neurons, with 5 neurons representing the familiar environment and 5 neurons representing the novel environment. The firing rate of the hippocampal neurons *r* is described by the following equation:

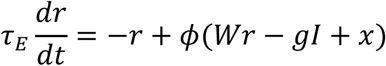

where *τ_E_* is the time constant of the neurons, *φ* is the activation function with rate clipping at *r_max_*, given by *φ*(*x*) = min(*r_max_*, max(0, *x*)), *W* is the recurrent weight matrix, *g* is the gain of the inhibitory input, *I* is the inhibitory feedback, and *x* is the external input. The inhibitory feedback is modeled as a fast global inhibition that reflects the overall activity of the network:

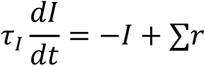

where *τ_I_* is the time constant of the inhibitory feedback. (hyperparameters: *dt*=0.001, *τ_E_*=0.1, *τ_I_*=0.02, *g*=0.3, *r_max_*=2.5)

The recurrent weight matrix *W* was initialized with 0 and updated according to the Hebbian-like learning rule in three separate phases, reflecting familiar environment, novel environment, and sleep after the novel environment. We modeled the sleep phase as a period of joint activity of the familiar and novel input patterns. During the learning phase, recurrent dynamics were turned off, and the network was clamped to the input pattern *x*. Input pattern *x* was 0 for inactive neurons, and was sampled from a Gaussian distribution with mean 1 and standard deviation 0.15 for active neurons.

The network was first exposed to the familiar environment, where the network was imposed with a familiar input pattern *x_F_* that activated only the familiar population. The weights were updated with a learning rate *η* for *t_F_* time steps.

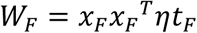

During the second phase, the network was exposed to a novel environment, where the network was imposed with a novel input pattern *x_N_* that activated only the novel population for *t_N_* time steps. The weights were updated similarly, giving

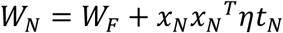

Finally, during the third phase, the network was exposed to a joint replay of the familiar and novel input patterns *x_R_* which activated all the neurons, and the weights were updated for *t_R_* time steps, giving

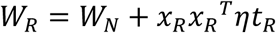

Weights were bounded to a maximum value *w_max_*=2.5 and self-weights were set to 0. The number of steps for each phase was set to *t_F_*=300, *t_N_*=30, *t_R_*=50 reflecting the relative duration of each phase. Learning rate was set to *η*=0.005.

For each phase, after the weight update, we simulated the recurrent dynamics of the network for 1000 time steps when it was given the novel input pattern *x_N_*.

## Statistical analysis

Statistical tests were performed in Python3.11 using SciPy (v1.15), Statsmodels (v0.14) and Scikit-Posthocs (v0.11), unless otherwise specified in figure legends. Depending on the dataset, Wilcoxon signed-rank test and Kruskal-Wallis test (followed by a post-hoc Dunn’s test) were applied as appropriate (information provided in the corresponding figure legends and result section). Linear regression was performed using ordinary least squares model from Statsmodels, reported R^2^ values were not adjusted for multiple regressors. The significance threshold α was set at 0.05. For multiple comparisons, p-values were adjusted using the Holm-Sidak method unless stated otherwise. Details of sample sizes n and p-values were provided in the corresponding figure legends. Data were presented as mean ± standard deviation in Results section and in figures, unless specified otherwise.

## Extended Data and Figures

**Extended Data Fig. 1:**
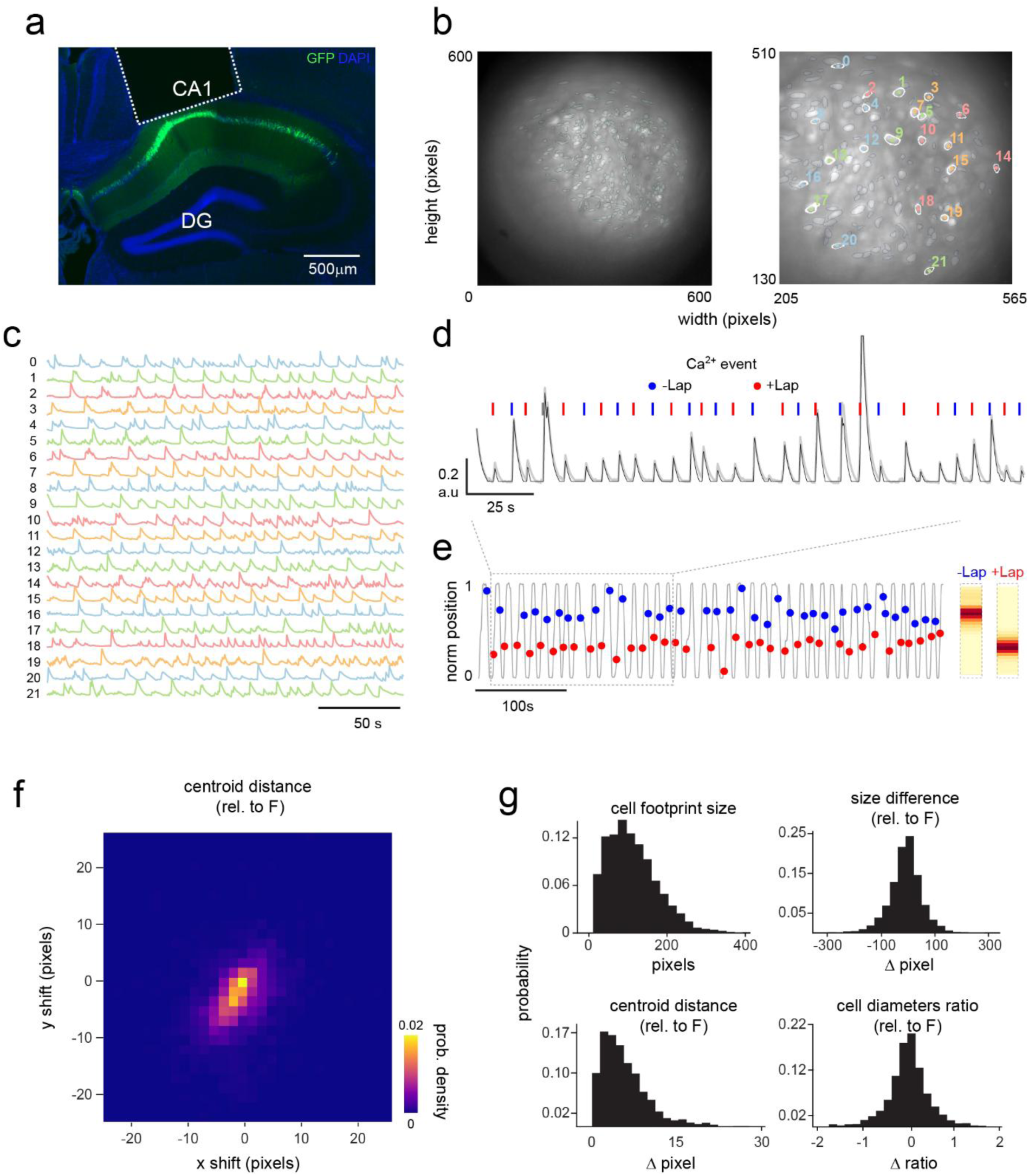
Miniscope Ca^2+^ imaging of CA1 place cells. (**a**) Coronal brain section showing viral expression of soma-targeted GCaMP7f in hippocampal CA1 of the right hemisphere. The position of the GRIN lens is shown as dotted line. Green: somaGCaMP7f; Blue: DAPI. Scale bar: 500 μm. DG: dentate gyrus. (**b**) Representative maximum-intensity projection of a CA1 recording acquired with the UCLA V4 Miniscope. **Left**: field of view (FOV) overlaid with the spatial footprints of all identified neurons (green outlines). **Right**: enlarged FOV showing 22 randomly selected place cells (white outlines; numbered). (**c**) Representative Ca^2+^ activity traces of the place cells shown in (b). Colors and numbers, same as in (b). (**d**) Representative Ca^2+^ activity trace from a place cell recording shown in (e) (gray trace: raw Ca^2+^ activity extracted by using CNMF-E algorithms, see Methods; black trace: smoothed Ca^2+^ activity). Colored ticks above trace indicate detected Ca^2+^ events during running in the positive (red) and negative (blue) lap directions. (**e**) Spatial activity of the same place cell shown in (d). Detected Ca^2+^ events are overlaid on the animal’s trajectory (gray). Red and blue dots indicate events during positive and negative running directions, respectively. Right: Gaussian-smoothed Ca^2+^ event rate maps for each running direction, normalized to the peak response (see Methods). (**f**) Distribution of centroid displacements of cross-registered cells relative to the reference (F environment). Color indicates probability density. Note that, in line with previous work^71^, absolute centroid positions shifted by 1.24±3.78 pixels in the horizontal direction and 2.97±4.67 pixels in the vertical direction (mean ± SD), corresponding to a mean centroid displacement of 5.57±3.93 pixels (see also in (g)). This displacement is smaller than the approximate cell radius, indicating accurate spatial alignment across environments (n_environment_=68; N_mice_=5). (**g**) Quantitative assessment of cross-registration accuracy across recordings based on somatic footprint area, centroid displacement, footprint size difference, and footprint shape indicating reliable alignment of individual cells across environments. **Top left**: distribution of cell footprint areas (111.59±63.22 pixels^2^, corresponding to a mean radius of 5.96 pixels; mean ± SD same for below). **Top right**: difference in footprint size relative to the reference environment (F) (8.77±59.31 pixels^2^). **Bottom left**: distribution of centroid displacements relative to F (5.57±3.93 pixels, see also in (f)). **Bottom right**: difference in the major-to-minor axis ratio of aligned footprints relative to F (0±0.44), indicating preservation of footprint shape across environments (n_environment_=68; N_mice_=5).

**Extended Data Fig. 2:**
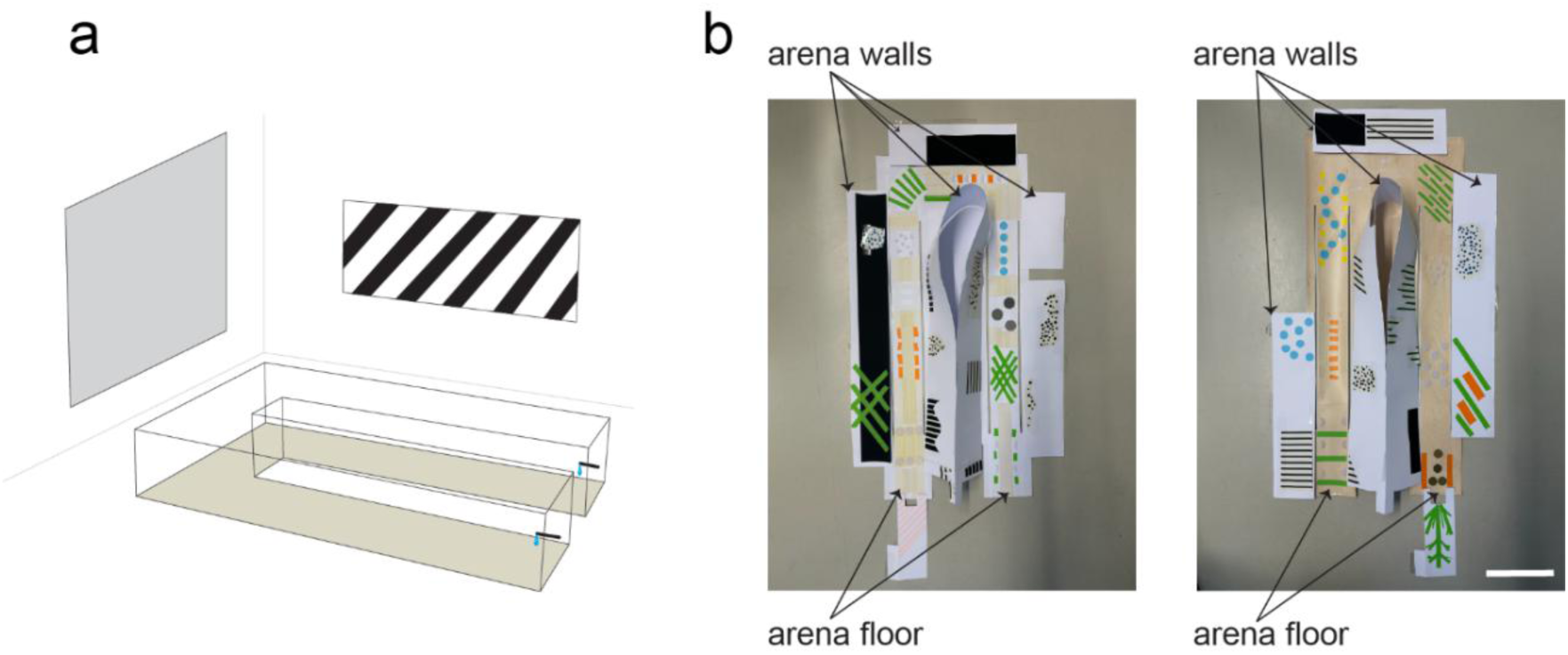
Local cues in familiar and novel environment. (**a**) Schematic of the experimental setup showing the U-shaped arena and distal cues. Distal cues were kept constant between F and N environments, whereas local cues on the arena floor and walls were changed (see b). (**b)** Representative photographs of the local cues in F and N environments. Scale bar: 14 cm.

**Extended Data Fig. 3:**
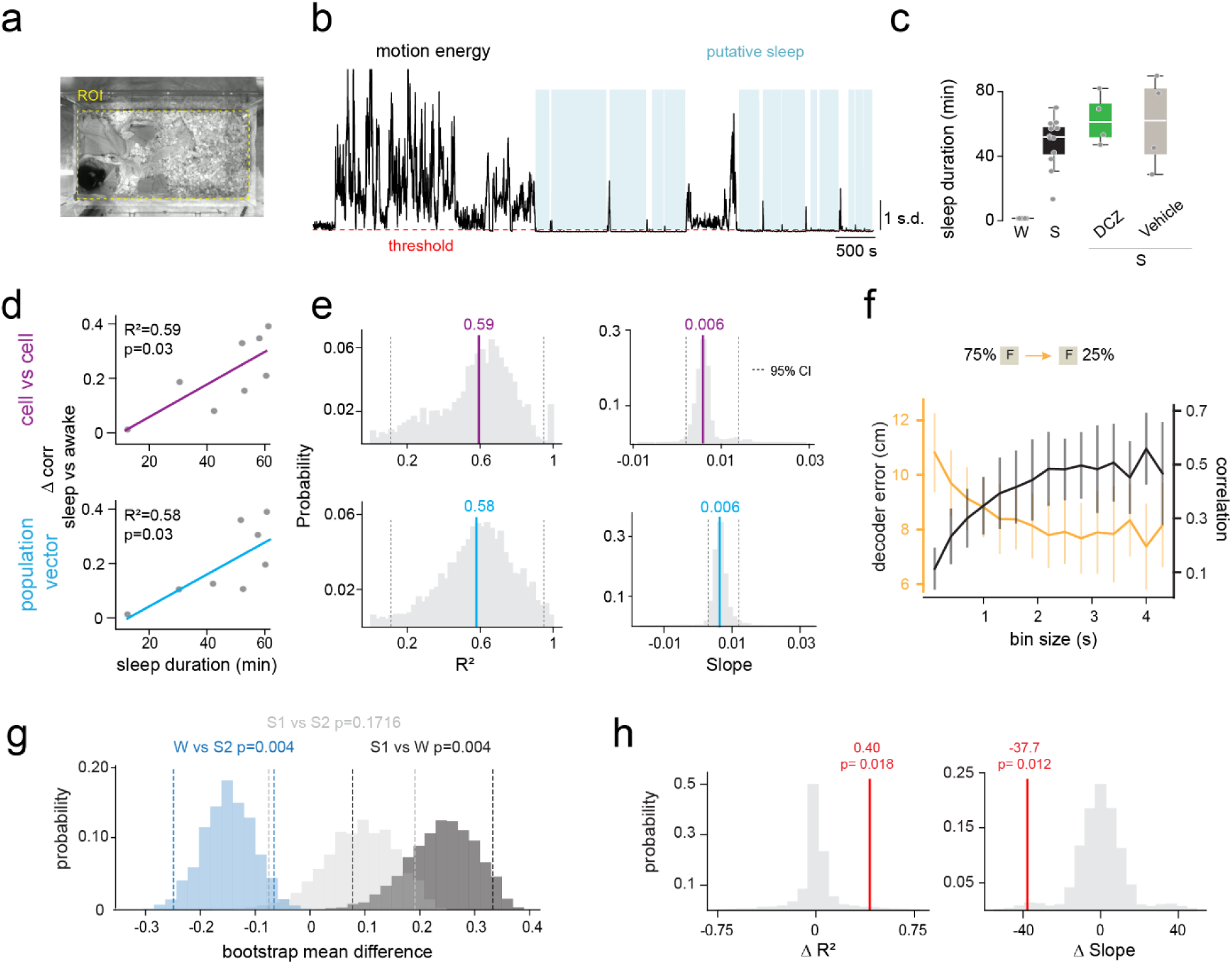
Videography-based sleep scoring and control analyses of decoding and regression across sleep and wake conditions. (**a**) Representative video frame of an animal during the post-encoding retention phase. Motion energy was quantified within the defined ROI (yellow rectangle). (**b**) Representative motion energy trace (black) from the same animal as in (a) showing putative sleep phases (blue). A conservative threshold corresponding to 0.1±0.05 of the standard deviation of the motion trace (red dashed line) was applied to identify immobile episodes (see Methods). Episodes with characteristic sleep posture (validated by visual inspection) and longer than 60 s were considered putative sleep episodes. The total sleep duration was computed as the sum of all validated sleep episodes. (**c**) Distribution of sleep duration across all trials under the wake (light gray), sleep (black), mPFC inactivation (DCZ, green) and vehicle (warm gray) conditions (wake: N_mice_=7, n_trials_=12; sleep: N_mice_=7, n_trials_=13; DCZ and vehicle: N_mice_=4, n_trials_=4). (**d**) Sleep vs. wake difference in F−N’ spatial rate map correlations positively correlated with total sleep duration, indicating longer sleep was associated with stronger generalization (Top: cell-by-cell correlations, R^2^=0.59; *p*=0.03; Bottom: PV correlations, R^2^=0.58; *p*=0.03; n_trials_=8, N_mice_=5) (**e)** Bootstrap analysis (5000 repetitions with replacement; n_trials_=8, N_mice_=5) assessing the robustness of the regression shown in (d). Observed regression parameters (R^2^ and slope) fell within the 95% bootstrap confidence intervals, with bootstrap confidence intervals for the slope remaining positive. (**f**) Decoder performance across different time bin sizes. Decoder error (left axis, orange) and Spearman correlation between decoded and true position (right axis, blue) were quantified by training on 75% of the place cell data in the familiar environment and testing on the remaining 25%. In line with a previous work^91^, the decoder accuracy changed as a function of the temporal bin size, i.e. the decoding error decreased and correlation between decoded and true position increased with larger bin sizes. A bin size at 2 s (see Methods) was chosen for subsequent analyses as it provided a good balance between accuracy and reliability. Error bar indicating mean ± SD. (**g**) Bootstrap analysis of condition-order effects (corresponding to Fig. 2c; resample size=4, 10^4^ repetitions; n_trials_=4, N_mice_=4). The distributions show bootstrapped mean differences in cell-by-cell correlation across conditions. Dashed lines represent the 95% confidence intervals. Significant differences were observed between sleep condition test 1 vs. wake (blue, S1 vs. W, *p*=0.004) and sleep condition test 2 vs. wake (dark gray, S2 vs. W, *p*=0.004), but not between the two sleep conditions (light gray, S1 vs. S2, *p*=0.172). (**h**) Permutation test of linear regression models shown in Fig. 3c (resample size=8, permutation of condition labels without replacement, 10^4^ repetitions; n_trials_=8, N_mice_=5). The observed difference in ΔR^2^ and Δslope values between sleep and awake conditions (red line) fell within the tail of the permutation distribution (ΔR^2^: *p*=0.018; Δslope: *p*=0.012), indicating a significant condition-dependent effect on regression fits.

**Extended Data Fig. 4:**
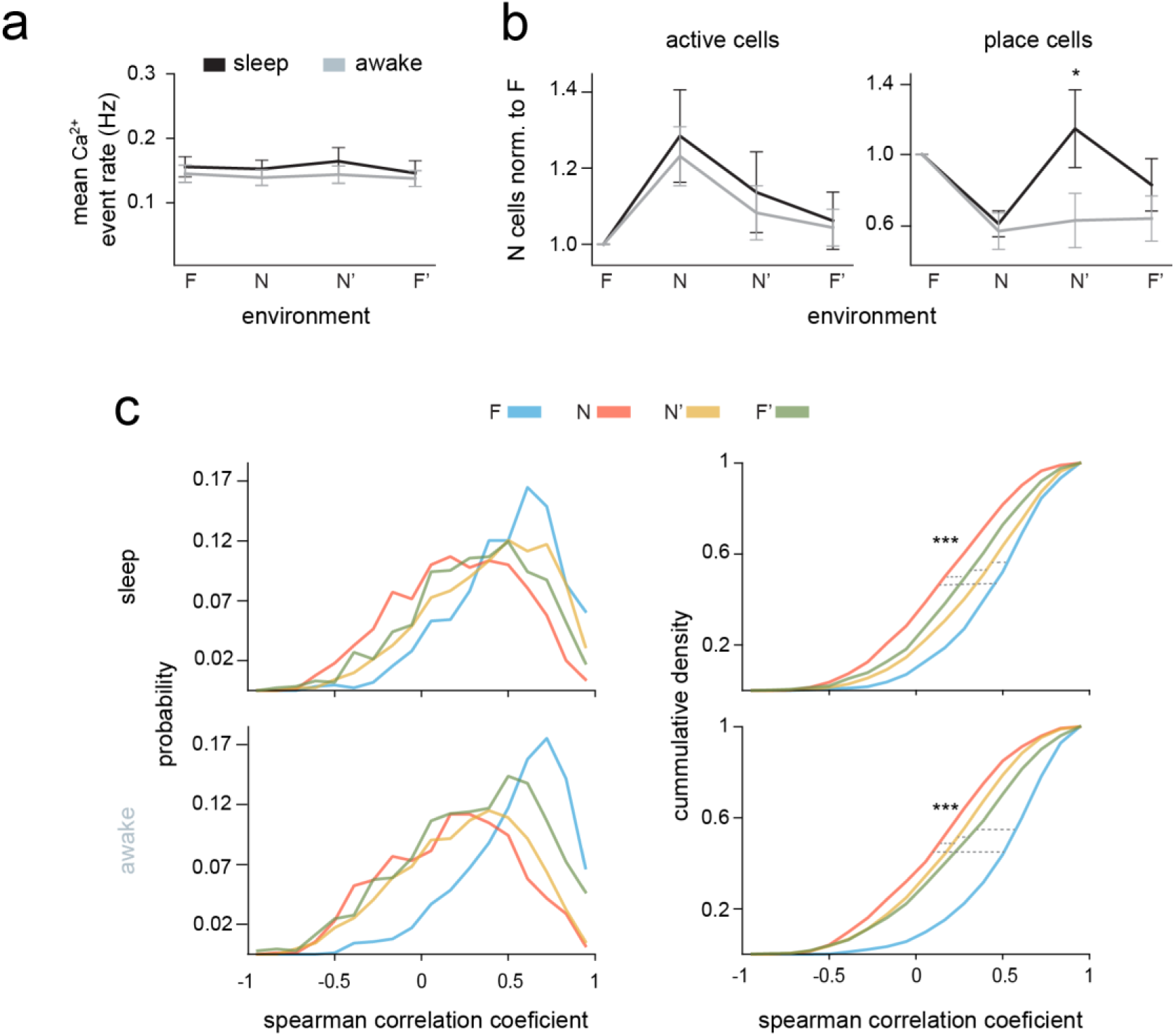
Activity and within-environment stability of CA1 neurons across environments and conditions. (**a**) Mean Ca^2+^ event rates for all active CA1 neurons during running across F, N, N’ and F’ under sleep (black) and awake (gray) conditions (9 sleep recordings and 8 awake recordings, in total n_environment_=68, N_mice_=5). Overall neuronal activity did not differ significantly across environments or conditions (Kruskal–Wallis test, *p*=0.064). Error bar indicating mean ± SD. (**b**) Relative proportion (relative to F) of all active cells (left) and place cells (right) across the F-N-N’-F’ recording conditions (as in Fig. 1a). In line with previous work^92–95^, the number of active cells increased as a result of novelty, both in sleep and wake condition, and no difference was found across conditions (Kruskal-Wallis test, *p*=0.247; n_environment_=68, N_mice_=5). However, the proportion of place cells in N’ returned to familiar like levels after sleep, but not after wake (Kruskal-Wallis test, *p*=0.002; Post-hoc Dunn’s test with Benjamini-Hochberg correction, *p*_(SF−SN)_=0.025, *p*_(SF−SN’)_=0.574, *p*_(WF−WN’)_=0.025, *p*_(SN’−WN’)_=0.024) in line with the generalization effect (see Fig. 2). Error bar indicating mean ± SD. (**c**) Within-environment stability of place cell representations. For each cross-registered place cell, spatial rate maps from the first and second halves of each recording (5 min each) were compared using Spearman’s correlation coefficients. Data are shown as probability density distributions (left) and cumulative distributions (right; same data). Note that sleep (top row) selectively increased within-environment stability in N’, whereas no comparable increase was observed following wake (bottom row). (two-sample Kolmogorov-Smirnov test with Bonferroni corrector, *p*<0.001 for all pairwise comparison; n_trials_=8, N_mice_=5).

